# Engineering *Saccharomyces cerevisiae* for growth on xylose using an oxidative pathway

**DOI:** 10.1101/2024.12.19.629315

**Authors:** Kenya Tanaka, Takahiro Yukawa, Takahiro Bamba, Miho Wakiya, Ryota Kumokita, Yong-Su Jin, Akihiko Kondo, Tomohisa Hasunuma

## Abstract

The fermentative production of valuable chemicals from lignocellulosic feedstocks has attracted considerable attention. Although *Saccharomyces cerevisiae* is a promising microbial host, it lacks the ability to efficiently metabolize xylose, a major component of lignocellulosic feedstocks. The xylose oxidative pathway offers advantages such as simplified metabolic regulation and fewer enzymatic steps. Specifically, the pathway involves the conversion of xylose into 2-keto-3-deoxy-xylonate, which can be channeled into two distinct pathways, the Dahms pathway and the Weimberg pathway. However, the growth of yeast on xylose as the sole carbon source through the xylose oxidative pathway has not been achieved, limiting its utilization. We successfully engineered *S. cerevisiae* to metabolize xylose as its sole carbon source via the xylose oxidative pathways, achieved by enhancing enzyme activities through iron metabolism engineering and rational enzyme selection. We found that increasing the supply of the iron-sulfur cluster to activate the bottleneck enzyme XylD by *BOL2* disruption and *tTYW1* overexpression facilitated the growth on xylose and the production of ethylene glycol at 1.5 g/L via the Dahms pathway. Furthermore, phylogenetic analysis of xylonate dehydratases led to the identification of a highly active homologous enzyme. A strain possessing the Dahms pathway with this highly active enzyme exhibited reduced xylonate accumulation. Furthermore, the introduction of enzymes based on phylogenetic tree analysis allowed for the utilization of xylose as the sole carbon source through the Weimberg pathway. This study highlights the potential of iron metabolism engineering and phylogenetic enzyme selection for the development of non-native metabolic pathways in yeast.

**Key points:**

- 1.5 g/L ethylene glycol was produced via Dahms pathway in *S. cerevisiae*.
- Enzyme activation enabled growth on xylose via both the Dahms and Weimberg pathways.
- Tested enzymes in this study may expand application of xylose oxidative pathway.

## Introduction

In recent decades, there has been an increase in interest on the fermentative production of high-value chemicals utilizing inedible lignocellulosic feedstocks, such as rice straw, wheat straw, bagasse residue after sugar cane juicing, corn stover, and switchgrass (Lane et al. 2018). Lignocellulosic feedstocks contain three main components: cellulose, hemicellulose, and lignin. Upon degradation, these feedstocks yield glucose, xylose, acetic acid, and formic acid. Organic acids, particularly acetic and formic acids, inhibit the growth of microorganisms, especially prokaryotes such as *Escherichia coli*, which is commonly used as a host for microbial chemical production (Lane et al. 2018). Conversely, the eukaryotic budding yeast *Saccharomyces cerevisiae* is considered to be a promising microbial host due to its ability to tolerate low pH and growth inhibitors (Jeffries 2006; Chen and Nielsen 2013; Liu et al. 2013). The weight ratio of the monosaccharide substrate obtained from lignocellulosic feedstocks is 30-50% glucose and 20-25% xylose (Kim et al. 2013). Therefore, for the direct production of compounds from the degradation products of lignocellulosic feedstocks, it is desirable to develop an engineered *S. cerevisiae* that is capable of utilizing both glucose and xylose.

There are two pathways for the utilization of xylose in yeast: the xylose reductase/xylitol dehydrogenase (XR/XDH) pathway and the xylose isomerase (XI) pathway. In the XR/XDH pathway, the enzymes xylose reductase (XR) and xylitol dehydrogenase (XDH) convert xylose to xylulose via xylitol. While XR has a dual cofactor preference with NADPH and NADH, XDH uses only NAD^+^, leading to cofactor imbalance during fermentation and low ethanol yields with accumulation of xylitol (Matsushika et al. 2009). On the other hand, xylose is converted directly into xylulose in the XI pathway, while conversion yields of isomerase reactions are limited by the inherent thermodynamic equilibrium between the substrate and product (Liu et al. 2019). Xylulose is then converted into xylulose-5-phosphate (X5P) by xylulose kinase (XK). X5P is metabolized via the yeast native pentose phosphate pathway and glycolysis. While the phosphorylation reaction by XK plays a crucial role in linking conventional xylose assimilation pathways with the endogenous pentose phosphate pathway, the activity of XK in *S. cerevisiae* needs improvement (Johansson et al. 2001). However, overexpression of *XKS1* from *S. cerevisiae* or XK from other microbes can lead to ATP depletion and inhibit yeast cell growth (Jin et al. 2003). Moreover, the pentose phosphate pathway, which is involved in nucleic acid synthesis, has a complex metabolic regulation mechanism (Masi et al. 2021). Nevertheless, recent studies have focused on reprogramming the regulatory networks, and improved xylose utilization through these pathways (Lee et al. 2021; Trivedi et al. 2023).

The xylose oxidative pathway offers another promising option to utilize xylose. This pathway utilizes non-phosphorylated and independent intermediates, thereby simplifying metabolic regulation and substrate competition when other carbon sources are present. The xylose oxidative pathway comprises thermodynamically advantageous reactions (Valdehuesa et al. 2018). In addition, the pathway involves fewer enzymatic reaction steps (Figure 1). Xylose is first oxidized to xylonolactone by NAD^+^-dependent xylose dehydrogenase XylB, and xylonolactone is converted into xylonate through a spontaneous ring-opening reaction or by the enzyme XylC. Xylonate is then converted into 2-keto-3-deoxy-xylonate (KDX) by the unique enzyme xylonate dehydratase XylD, which contains an iron-sulfur cluster. KDX serves as a crucial branching point, leading to two distinct metabolic pathways. In the Dahms pathway, KDX is converted into pyruvate and glycolaldehyde by the KDX aldolases YihH or YagE. Pyruvate is metabolized via the TCA cycle and glycolaldehyde is further converted into its oxidative form, glycolate, or its reductive form, ethylene glycol. In the Weimberg pathway, KDX is converted into α-ketoglutarate semialdehyde (AKGSA) by 2-keto-3-deoxy-xylonate dehydratase XylX. Finally, AKGSA is converted into α-ketoglutarate (AKG) by AKGSA dehydrogenase. Therefore, the xylose oxidative pathway requires fewer than five steps to synthesize important metabolites such as pyruvate or α-ketoglutarate, which are then metabolized via glycolysis and the TCA cycle, leading to cell growth and bioproduction. Thus, the xylose oxidative pathway is a shortcut from xylose to hub-intermediates, facilitating the production of useful compounds from glucose and xylose.

**Figure 1.**
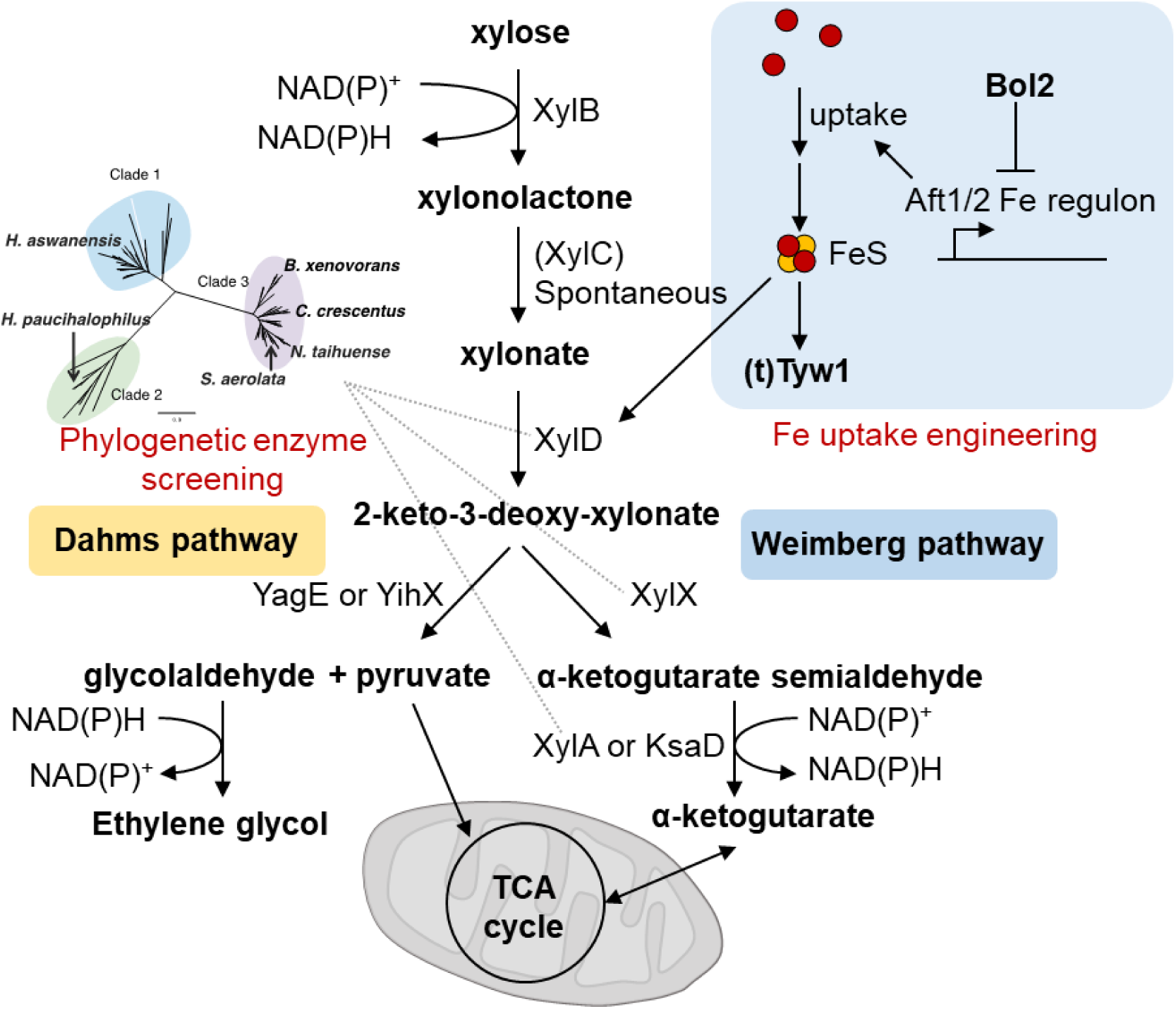
Construction of xylose oxidative pathway based on enzyme activation by iron uptake engineering and phylogenetic enzyme screening. XylB; Xylose dehydrogenase, XylD; Xylonate dehydratase, XylX; 2-keto-3-deoxy-xylonate dehydratase, KsaD; 2-ketoglutarate semialdehyde dehydrogenase, YagE; 2-keto-3-deoxy-xylonate aldolase.

The gene cluster of the *xylXABCD* operon in the Weimberg pathway has been extensively characterized in *Caulobacter crescentus* (Stephens et al. 2007; Watanabe et al. 2006; Almqvist et al. 2018). This operon enables *Pseudomonas putida* to grow on xylose as the sole carbon source with a specific growth rate of 0.21 h^−1^ (Meijnen et al. 2009). Subsequently, it was discovered that *Escherichia coli* and *Corynebacterium glutamicum* can also grow on xylose using the Weimberg pathway (Rossoni et al. 2018; Tai et al. 2016; Brüsseler et al. 2019). Moreover, *E. coli* has been engineered to produce both mesaconic acid, poly(lactate-co-glycolate), and itaconic acid via the xylose oxidative pathway (Lu et al. 2021; Choi et al. 2016; Bai et al. 2016). A pathway design termed “parallel metabolic pathway engineering” using the xylose oxidative pathway was shown to be effective for co-utilization of glucose and xylose (Fujiwara et al. 2020). *Bacillus subtilis* can also produce polyglutamic acid from xylose via the Weimberg pathway (Halmschlag et al. 2020). However, while attempts have been made to cultivate the yeast *S. cerevisiae* on xylose as the sole carbon source via the xylose oxidative pathway (Borgström et al. 2019; Wasserstrom et al. 2018), this has not yet been achieved. One possible reason for this is the difficulty associated with expressing enzymes with iron-sulfur clusters, such as XylD, in an active state, as this is a bottleneck reaction in yeast. To address this issue, XylD activation by *BOL2* destruction has been performed to force *S. cerevisiae* to use either the Dahms pathway or the Weimberg pathway (Borgström et al. 2019; Salusjärvi et al 2017). A previous study also showed that *BOL2* disruption and t*TYW1* overexpression can improve XylD activity in *S. cerevisiae* (Bamba et al. 2019).

Another problem in engineering the xylose oxidative pathway in *S. cerevisiae* is the limited availability of enzymes that offer differential catalytic activities. Conventionally, enzymes from *C. crescentus* have been used, following the identification of the *xylXABCD* gene operon associated with the Weimberg pathway in *C. crescentus* in 2007 (Stephens et al. 2007). Despite multiple studies investigating enzymes from taxa other than *C. crescentus*, highly active enzymes enabling growth on xylose as the sole carbon source have not yet been identified in yeast. Therefore, a systematic classification and comparison of enzyme activities from various sources is needed for the construction of a functional xylose oxidative pathway in yeast.

In this study, we sought to culture *S. cerevisiae* using xylose as the sole carbon source via oxidative xylose metabolic pathways. By disrupting *BOL2* and overexpressing *tTYW1*, we enhanced the supply of iron-sulfur clusters for XylD, enabling xylose-dependent growth through the Dahms pathway. In addition, by introducing highly active enzymes such as XylX from *Burkholderia xenovorans, Sphingomonas aerolata,* or *Novosphingobium taihuense*, we were able to induce xylose-dependent growth using the Weimberg pathway. Our results demonstrate the potential of using enzymes from various organisms for constructing a xylose oxidative pathway in yeast, and suggest that a systematic classification approach can be useful for identifying highly active enzymes. Our findings have important implications for the development of yeast as a platform for producing value-added chemicals from renewable sources, such as lignocellulosic feedstocks.

## Materials and Methods

### Constructing plasmids

The plasmids and primers used in this study are shown in Supplemental Tables S1 and S2. *E. coli* NovaBlue (Merck Millipore, Darmstadt, Germany) was used for plasmid amplification. NovaBlue was cultured in Luria-Bertani medium (10 g/L tryptone, 5 g/L yeast extract, and 5 g/L NaCl) containing 100 μg/mL ampicillin at 37 °C. Cloning was performed using the In-Fusion® HD Cloning Kit (Takara Bio, Shiga, Japan) and Ligation High Ver. 2 (Takara Bio). Briefly, the homologous sequence of the terminal 15 bases of the restriction enzyme-treated vector was added to both ends of the primers to be cloned by PCR. The vector, the PCR product, and in-fusion enzyme were mixed and then incubated at 50 °C for 15 minutes. All of the genes used in this study were artificially synthesized by GenScript (Piscataway, NJ, US) with DNA sequences optimized for the codon usage frequency of *S. cerevisiae*. The nucleotide sequences are shown in Supplemental Tables S3.

To introduce the Dahms pathway into yeast, YagE from *E. coli* was artificially synthesized and cloned into pIL-pTDH3-tADH1 using *Sma*I, following the method described by Bamba et al. (2019). By cloning the *ksaD* gene from *C. glutamicum*, we generated the plasmid pIU-pTDH3-CgksaD. The SED1 promoter region and SAG1 terminator were obtained by PCR using YPH499 genomic DNA as a template with specific primers (xhoI-SED1p F, xhoI-SED1p R, SAG1t F, and SAG1t R). The resulting DNA fragments were treated with *Xho*I and *Not*I and cloned into pGK405 (LEU2) using in-fusion cloning, resulting in the creation of pIL-pSED1-tSAG1 (Ishii et al. 2009).

Next, pIL-pSED1-tSAG1 was treated with *Sma*I, and *xylX* genes from *N. taihuense*, *H. aswanensis*, *H. paucihalophilus*, *C. crescentus*, *B. xenovorans*, and *S. aerolata* were artificially synthesized and cloned into pIL-pSED1-tSAG1, resulting in the construction of pIL-pSED1-HaxylX-tSAG1, pIL-pSED1-HpxylX-tSAG1, pIL-pSED1-CcxylX-tSAG1, pIL-pSED1-BxxylX-tSAG1, pIL-pSED1-SαxylX-tSAG1, and pIL-pSED1-NtxylX-tSAG1, respectively. To construct pIA-pTDH3-tADH1, pIL-pTDH3p-tADH1 and pGK402 (ADE2) (Ishii et al. 2009) were cloned by ligation. *xylB* from *C. crescentus* was artificially synthesized and cloned into *Sma*I-treated pIA-pTDH3-tADH1, resulting in the creation of pIA-pTDH3-xylB.

Artificial synthesis of *xylA* genes from various organisms, including *B. xenovorans*, *Asticcacaulis endophyticus*, *Streptomyces hygroscopicus*, *Pseudomonas inefficax*, *Bacteroidetes bacterium*, *Rhodopirellula sp. TMED11*, *Pseudomonas fluorescens*, *Sphingomonas aurantiaca*, *Rhodobacteraceae bacterium*, and *Burkholderia lata*, was performed. These genes were cloned into pIU-pTDH3-tADH1 using *Sma*I, resulting in the creation of pIU-pTDH3-BxxylA, pIU-pTDH3-AexylA, pIU-pTDH3-ShxylA, pIU-pTDH3-PixylA, pIU-pTDH3-BbxylA, pIU-pTDH3-RspxylA, pIU-pTDH3-PfxylA, pIU-pTDH3-SaxylA, pIU-pTDH3-RbxylA, and pIU-pTDH3-BlxylA.

To create plasmids expressing *xylD* from heterologous species, pGK406 (URA3) (Ishii et al. 2009) and pIL-pSED1-tSAG1 were digested with *Xho*I and *Not*I, and then ligated to create pIU-pSED1-MaxylD, pIU-pSED1-CcxylD, pIU-pSED1-BxxylD, pIU-pSED1-HexylD, pIU-pSED1-SexylD, pIU-pSED1-AtxylD, pIU-pSED1-RmxylD, and pIU-pSED1-PaxylD, using artificially synthesized DNA from *Marinovum algicola*, *Caulobacter crescentus*, *Burkholderia xenovorans*, *Herbaspirillum hiltneri*, *Sphingomonas elodea*, *Agrobacterium tumefaciens*, *Rhizobium miluonense*, and *Pseudooceanicola antarcticus*, respectively.

### Construction of yeast strains

The yeast strains constructed in this study are shown in Supplemental Table S4. Yeast transformation was performed using a one-step transformation method with lithium acetate (Chen et al., 1992). *Saccharomyces cerevisiae* YPH499 [MATa ura3-52 lys2-801 ade2-101 trp1-63 his3-Δ200 leu2-Δ1 (Stratagene, La Jolla, CA, USA)] was used to construct the oxidative xylose metabolic pathway. The yeast strains were cultured in SD medium (6.7 g/L yeast nitrogen base without amino acids (Difco Laboratories, Detroit, MI, USA), 20 g/L glucose) with amino acids and nucleic acids added to meet the nutritional requirements of the strain. SCD medium (6.7 g/L yeast nitrogen base without amino acids (Difco Laboratories), 2 g/L synthetic complete mix, 20 g/L glucose) was also used. During yeast transformation, amino acids and nucleic acids were excluded from the synthetic complete mix based on the nutritional requirements of the strains.

The BD strain was constructed by introducing the *xylB* and *xylD* genes from *C. crescentus* and disrupting *GRE3*, which converts xylose to xylitol and xylonic acid, into *S. cerevisiae* YPH499. The BDE strain was generated by introducing pIL-pTDH3-yagE into the BD strain to incorporate the Dahms pathway. The BDΔB-E strain was created by introducing pIL-pTDH3-yagE and replacing the *BOL2* gene of the BD strain with *HIS3*. To disrupt the *BOL2* gene, DNA fragments containing the *ADE2* marker were obtained by performing PCR using pGK402 (ADE2) (Ishii et al. 2009) as the template and the primers dBOL2_ADE2 F and dBOL2_ADE2 R. PCR was also performed using *S. cerevisiae* genomic DNA as the template and the primers BOL2 up F, BOL2 up R, BOL2 down F, and BOL2 down R to amplify the 500-bp regions upstream and downstream of the BOL2 ORF. The BOL2 disruption fragments, comprising the ADE2 marker flanked by the 500 bp ORF regions, were generated by overlapping PCR and subsequently transformed into the BD strain. The BDΔBtT strain was generated by introducing pIAur-tTYW1 into the BDΔB strain. The BDΔBtT-E strain and BDΔBtT-H strains were generated by integrating pIL-pTDH3-yagE and pIL-pTDH3-yjhH, respectively, into the BDΔBtT strain to incorporate the Dahms pathway. To examine *xylD* activity in heterologous organisms, the base BE strain was generated by introducing pIA-pTDH3-xylB and pIL-pTDH3-yagE into the YPH499ΔGRE3 strain. Subsequently, the BEΔBtT strain was obtained by replacing the *BOL2* gene of the BE strain with *HIS3*, and further introducing pIAur-tTYW1. Using the BEΔBtT strain as a base, the plasmids pIU-pSED1-MaxylD, pIU-pSED1-CcxylD, pIU-pSED1-BxxylD, pIU-pSED1-HexylD, pIU-pSED1-SexylD, pIU-pSED1-AtxylD, pIU-pSED1-RmxylD, and pIU-pSED1-PaxylD were transformed into it, resulting in the creation of BEΔBtT-MaxylD, BEΔBtT-CcxylD, BEΔBtT-BxxylD, BEΔBtT-HexylD, BEΔBtT-SexylD, BEΔBtT-AtxylD, BEΔBtT-RmxylD, and BEΔBtT-PaxylD strains, respectively.

To introduce the Weimberg pathway into the BDΔBtT strain, we first generated the BDΔBtT-ksaD strain by introducing pIU-pTDH3-CgksaD. Subsequently, strains such as BDΔBtT-ksaD-CcxylX, BDΔBtT-ksaD-HaxylX, BDΔBtT-ksaD-HpxylX, BDΔBtT-ksaD-BxxylX, BDΔBtT-ksaD-SaxylX, and BDΔBtT-ksaD-NtxylX were created by transforming pIL-pSED1-CcxylX-tSAG1, pIL-pSED1-HaxylX-tSAG1, pIL-pSED1-HpxylX-tSAG1, pIL-pSED1-BxxylX-tSAG1, pIL-pSED1-SaxylX-tSAG1, and pIL-pSED1-NtxylX-tSAG1, respectively.

To introduce *xylA* from heterologous organisms into yeast, BDΔBtT-BxxylX strains were created by introducing pIL-pSED1-BxxylX-tSAG1 into the BDΔBtT strain. In addition, by introducing pIU-pTDH3-BxxylA, pIU-pTDH3-AexylA, pIU-pTDH3-ShxylA, pIU-pTDH3-PixylA, pIU-pTDH3-BbxylA, pIU-pTDH3-RsxylA, pIU-pTDH3-PfxylA, pIU-pTDH3-SaxylA, pIU-pTDH3-RbxylA, and pIU-pTDH3-BlxylA, strains, BDΔBtT-BxxylX-BxxylA, BDΔBtT-BxxylX-AexylA, BDΔBtT-BxxylX-ShxylA, BDΔBtT-BxxylX-PixylA, BDΔBtT-BxxylX-BbxylA, BDΔBtT-BxxylX-RspxylA, BDΔBtT-BxxylX-PfxylA, BDΔBtT-BxxylX-SaxylA, BDΔBtT-BxxylX-RbxylA, and BDΔBtT-BxxylX-BlxylA were obtained, respectively.

### Growth test using xylose as a single carbon source

Cells were cultured in SD medium at 30 °C with gentle shaking at 200 rpm for 24 hours. The cultured cells were then washed twice with distilled water. After washing, the cells were inoculated to a 200-mL Erlenmeyer flask containing 50 mL of growth test medium consisting of 6.7 g/L yeast nitrogen base without amino acids (Difco Laboratories), 1.46 g/L Yeast Synthetic Drop-out Media Supplements (Y1501, Sigma Aldrich, Burlington, United States), 76 mg/L uracil, and 20 g/L xylose. Cells were inoculated at an optical density of OD_600_ = 0.1 and cultured at 30 °C with gentle shaking at 200 rpm. Cell density was measured using a UV-VIS spectrophotometer (uVmini-1240, Shimadzu Corp. Kyoto, Japan).

### Culture test with mixed sugar using glucose and xylose

Cells were cultured in 5 mL of SCD medium for 24 hours. Then, they were transferred to a 100-mL Erlenmeyer flask containing 20 mL of YPD medium with an optical density of OD_600_ = 0.1 and cultured at 30 °C with gentle shaking at 150 rpm for 24 hours. After culturing, the cells were washed twice with distilled water. They were then inoculated into 100-mL Erlenmeyer flasks containing YPDX medium (containing 10 g/L yeast extract, 20 g/L tryptone, 10 g/L glucose, and 10 g/L xylose) with an optical density of OD_600_ = 5.0. Cultures were performed at 30 °C with gentle shaking at 200 rpm in a total volume of 20 mL.

### Phylogenetic tree analysis

Amino acid sequences of the enzymes for phylogenetic tree analysis were retrieved from the NCBI database. Sequences that overlapped and those containing unknown amino acids were excluded from the analysis. In addition, CD-HIT (Fu et al. 2012) was used to remove amino acid sequences exhibiting more than 90% identity. Alignment was performed using MAFFT (Katoh et al. 2002) and phylogenetic trees were constructed with RaxML, based on the aligned sequence (Stamatakis et al. 2014).

### XylX activity measurement

Yeast strains were cultured for 24 hours at 30 °C in a 100-mL Erlenmeyer flask containing 20 mL of YPD medium (containing 10 g/L yeast extract, 20 g/L peptone, and 20 g/L glucose). The cells were then washed twice with wash buffer (10 mM phosphate buffer, 2 mM EDTA, pH = 7.5) and then exposed to protein extraction buffer (100 mM phosphate buffer, pH = 7.5, 2 mM MgCl_2_, 1 mM DTT) containing 200 μL of 0.5-mm diameter glass beads (YGB05) (Yasui Kikai, Osaka, Japan). The mixture was shaken at 1500 rpm for 5 minutes using a Shake Master Neo (Biomedical Science, Tokyo, Japan), followed by centrifugation at 21,000 × *g* for 15 minutes at 4 °C. The supernatant was used as the yeast total protein solution for enzyme activity measurement.

The following substrates were used for the XylX activity assay: 5 mM 2-keto-3-deoxy-xylonate (KDX), 20 mM Tris-HCl (pH = 8.0), 10 mM MgCl_2_, and 1 mM NAD+. A total of 50 μL of yeast protein extract was added to 350 μL of the substrate mixture and incubated at 30 °C for 5 minutes. To calculate the XylX activity, the increase in NADH during the reaction was measured over time at a wavelength of 340 nm. The molar absorption coefficient of NADH (6.3 mM^−1^ cm^−1^) was used to calculate the converted amount of KDX.

### XylD activity measurement

The activity of xylonate dehydratase (XylD) was measured by the thiobarbiturate method (Kim and Lee 2005). All proteins were extracted using the same method employed for the measurement of XylX activity. For the reaction solution, 400 μL of reaction mixture containing 50 mM Tris– HCl buffer (pH = 8.0), 5 mM MgCl_2_, and 12 mM D-xylonate, and 50 μL of yeast protein extraction solution were incubated at 30 °C for 10 minutes. After 10 minutes, the enzyme reaction was stopped by adding 100 μL of 2 M HCl. Subsequently, 50 μL of the solution was mixed with 125 μL of 25 mM iodic acid solution (dissolved in 0.125 M H_2_SO_4_) and reacted at 20 °C for 20 minutes. The reaction was then stopped by adding 250 μL of 2% (w/v) sodium arsenite solution (dissolved in 0.5 M HCl). Finally, 1 mL of 0.3% aqueous thiobarbiturate solution was added and incubated at 100 °C for 10 minutes to generate a red chromophore. To enhance the color intensity of the red chromophore, an equal amount of DMSO was added to the reaction solution cooled to room temperature. The specific activity of XylD was calculated based on the absorbance at 549 nm. The molar absorption coefficient of the red chromophore is 6.78 × 10^4^ M^−1^ cm^−1^ (Skoza and Mohos 1976).

### Metabolite analysis

Metabolite concentrations were measured as described in Bamba et al. (2019). Briefly, fermentation supernatants (5 μL) containing 10 g/L ribitol as an internal standard (2 μL) were evaporated to dryness using a CentriVap Benchtop Vacuum Concentrator (Labconco, Kansas City, MO, USA). Sample derivatization was performed in a shaker incubator (1200 rpm at 30 °C; MBR022UP; Taitec, Saitama, Japan) for 90 minutes with 100 μL of a 20 mg/mL solution of methoxyamine hydrochloride in pyridine. This was followed by a 30-minute reaction in a shaker incubator (1200 rpm at 37 °C; M-BR-022UP; Taitec) with 50 μL of N-methyl-n-TMS-trifluoroacetamide (MSTFA). After centrifugation at 3000 × *g* for 5 minutes at room temperature, 120 μL of the supernatant was transferred to a 150 μL glass insert and analyzed by GC-MS (GCMS-QP 2010 Ultra; Shimadzu Corp.) equipped with a CP-Sil 8CB column (30 m length × 0.25 mm i.d., 0.25 μm layer thickness; Agilent Technologies, Palo Alto, CA, USA). The operating parameters of the GC-MS were the same as those described previously (Bamba et al. 2019).

## Results

### Functional Construction of the Dahms Pathway

In *S. cerevisiae*, xylose is converted into xylitol by the endogenous non-specific aldose reductase encoded by the *GRE3* gene. To enhance the conversion efficiency of xylose via the oxidative xylose metabolic pathways, *GRE3* was disrupted in the YPH499 strain by replacing with the KanMX cassette, producing the base strain YPH499ΔGRE3.

To incorporate the Dahms pathway into yeast, the *yagE* gene from *E. coli* was introduced into the BD strain. This strain was constructed by integrating the *xylB* and *xylD* genes from *C. crescentus* into the YPH499ΔGRE3 strain (Bamba et al., 2019). However, when the growth of the resulting BDE strain was evaluated after culture in a medium containing xylose as the sole carbon source, it did not exhibit an increase in growth compared to the YPH499 wild type strain (Figure 2). We hypothesized that this was due to the limited activity of XylD caused by a deficiency of iron-sulfur clusters in yeast. Consequently, our objective shifted towards expressing an active form of XylD, complete with its iron-sulfur cluster, to facilitate the growth of the BDE strain on xylose by applying iron uptake engineering. Iron uptake in yeast involves the regulatory actions of Bol2 on the transcription factors Aft1/Aft2, which control iron uptake (Kumánovics et al. 2008; Courel et al. 2005; Kaplan and Kaplan 2009). When the *BOL2* gene was disrupted in the BDE strain, the resulting BDΔB-E strain demonstrated the capacity to grow on xylose (Figure 2), although the specific growth rate of BDΔB-E strain (*`μ`* = 0.035 h^−1^) is pretty low compared to that of a XI pathway strain (μ = 0.24 h^−1^) (Trivedi et al. 2023). The Tyw1 protein is hypothesized to bind excess Fe-S clusters, and the overexpression of a truncated form of Tyw1 (tTyw1) is known to enhance the expression of Aft1/Aft2 Fe regulon genes (Li et al 2011). Incorporating overexpressed tTyw1 into the BDΔB-E strain, resulting in the BDΔBtT-E variant, further improved growth on xylose. Conversely, loss of the *yagE* gene from the BDΔBtT-E strain impaired growth (Figure 2; BDΔBtT-empty strain). These results suggest that the engineering of iron uptake effectively resolved the issue of limited activity of XylD, thereby enabling the engineered yeast strains to grow on xylose by using the Dahms pathway although the growth rate remains to be improved.

**Figure 2.**
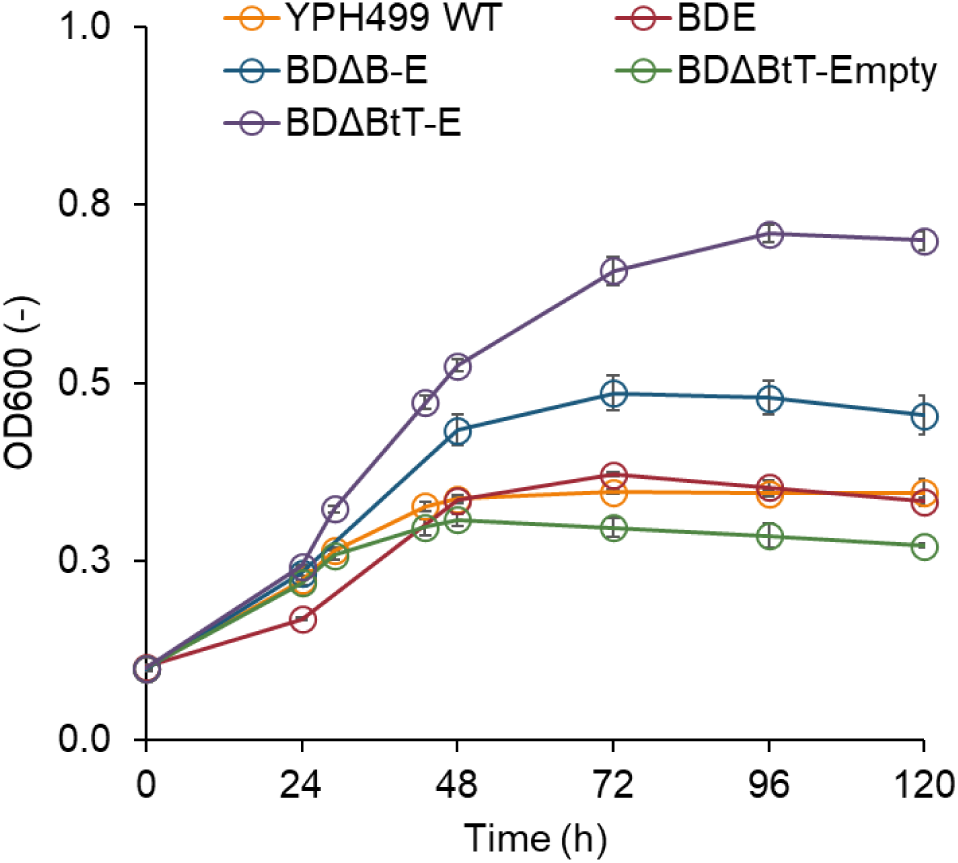
Growth on xylose as the sole carbon source by the Dahms pathway. Growth tests were performed on a medium containing 20 g/L of xylose. Values are the mean ± SD of three biological replicates.

In the context of the practical fermentation of a lignocellulosic hydrolysate, both glucose and xylose need to be metabolized. Therefore, we investigated whether the ability to utilize xylose via the Dahms pathway can enhance growth in a culture medium containing glucose and xylose. Under mixed sugar conditions, the BDΔBtT-E strain exhibited better growth than YPH499ΔGRE3 (Figure 3A). Furthermore, to assess the efficacy of different KDX aldolases, we introduced the *yjhH* from *E. coli*, replacing *yagE* to create the BDΔBtT-H strain. This modification also showed improved growth relative to YPH499ΔGRE3 (Figure 3A). While YPH499ΔGRE3 did not consume xylose, both the BDΔBtT-E and BDΔBtT-H strains completely utilized xylose (Figure 3C). However, the accumulation of approximately 8 g/L xylonate indicated suboptimal xylose utilization efficiency (Figure 3D). Nonetheless, both BDΔBtT-E and BDΔBtT-H strains produced 1.5 g/L of ethylene glycol (Figure 3E), a yield exceeding those reported in prior studies which produced ethylene glycol via the same pathway in *S. cerevisiae* (Salusjärvi et al. 2017). The similar performance of BDΔBtT-E and BDΔBtT-H strains suggests that *yagE* and *yjhH* are functionally equivalent in terms of constructing the Dahms pathway.

**Figure 3.**
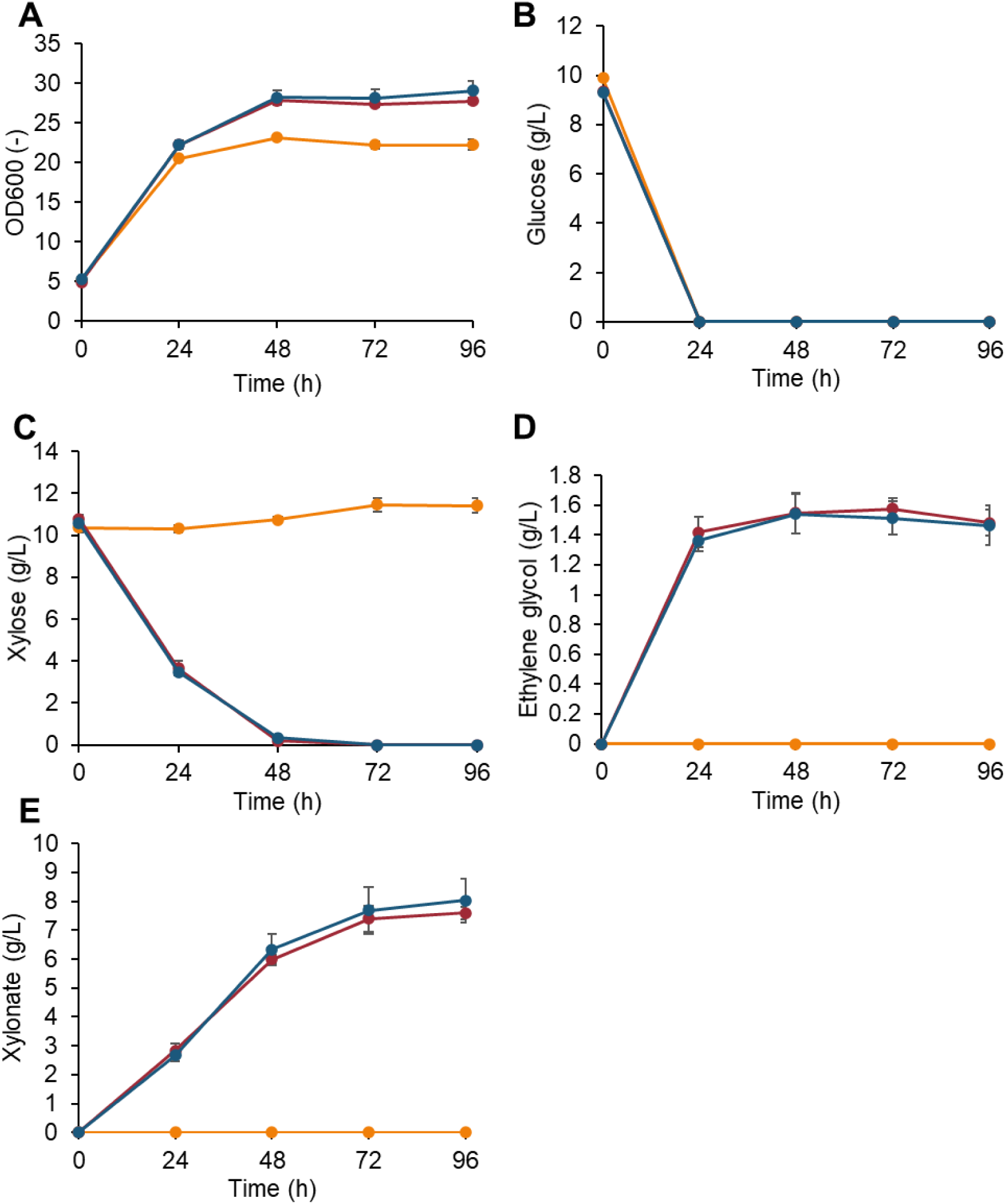
Mixed sugar culture of yeast strains with the Dahms pathway and enhanced Fe uptake systems. YPH499ΔGRE3 (yellow line), BDΔBtT-H (red line), and BDΔBtT-E (blue line) strains grown in YPDX medium (containing 10 g/L yeast extract, 20 g/L tryptone, 10 g/L glucose, and 10 g/L xylose). OD_600_ (A), substrates (B,C) and metabolites (D,E) were measured. Values are the mean ± SD of three biological replicates.

### Identification of highly active XylDs through phylogenetic analysis

Despite employing iron uptake engineering to augment the availability of iron-sulfur clusters for XylD, the observed accumulation of xylonate accumulation (∼8 g/L) suggests that the xylonate dehydratase activity is still limited. A previous study introduced XylD and YjhG from *Burkholderia cenocepacia*, *Haloferax volcanii*, *Ellin329* bacterium, and *E. coli* into *S. cerevisiae*, but these enzymes did not show improved cell growth or compound productivity compared to XylD from *C. crescentus* (Wasserstrom et al. 2018). As approximately 6,000 amino acid sequences are registered as XylD in NCBI, there may be XylDs with higher activities than that in *C. crescentus*.

Phylogenetic tree analysis is effective for identifying highly active enzymes for metabolic engineering (Protzko et al. 2018; Vavricka et al. 2019). Based on the phylogenetic tree analysis, xylonate dehydratases belonging to the IlvD/EDD family were classified into six clades (Figure 4A). We measured the xylonate dehydratase activity of six XylDs from *Marinovum algicola*, *Caulobacter crescentus*, *Burkholderia xenovorans*, *Herbaspirillum hiltneri*, *Sphingomonas elodea*, and *Agrobacterium tumefaciens*. Notably, four XylDs from *B. xenovorans*, *S. elodea*, *H. hiltneri*, and *A. tumefaciens* showed 1.85-, 9.62-, 3.46- and 23.9-fold higher activities, respectively, than that from *C. crescentus*, which has conventionally been used for constructing the xylose oxidative pathway (Figure 4B). To identify XylD with even higher activities than the abovementioned, we selected and measured the activity of two additional XylDs from *Rhizobium miluonense* and *Pseudooceanicola antarcticus*, which belong to the same clade as *A. tumefaciens*, whose XylD showed the highest activity in the previous selection round via phylogenetic tree analysis. Interestingly, XylD from *R. miluonense* and *P. antarcticus* showed 29.2-fold and 30.8-fold higher activities compared to that from *C. crescentus* (Figure 4B). Therefore, we constructed engineered yeast strains (BEΔBtT-RmXylD and BEΔBtT-PaXylD) harboring the Dahms pathway with the highly active XylDs (RmXylD and PaXylD) and performed mixed sugar cultivation. As a result, the accumulation of xylonate in BEΔBtT-PaXylD slightly decreased compared to the BEΔBtT-CcXylD strain (Figure 4C). However, the decreased xylonate accumulation in the BEΔBtT-PaXylD strain did not improve ethylene glycol production (Figure 4D). This may be due to increased generation of byproducts other than ethylene glycol. The BEΔBtT-RmXylD strain showed similar xylonate accumulation and ethylene glycol production to the BEΔBtT-CcXylD strain.

**Figure 4.**
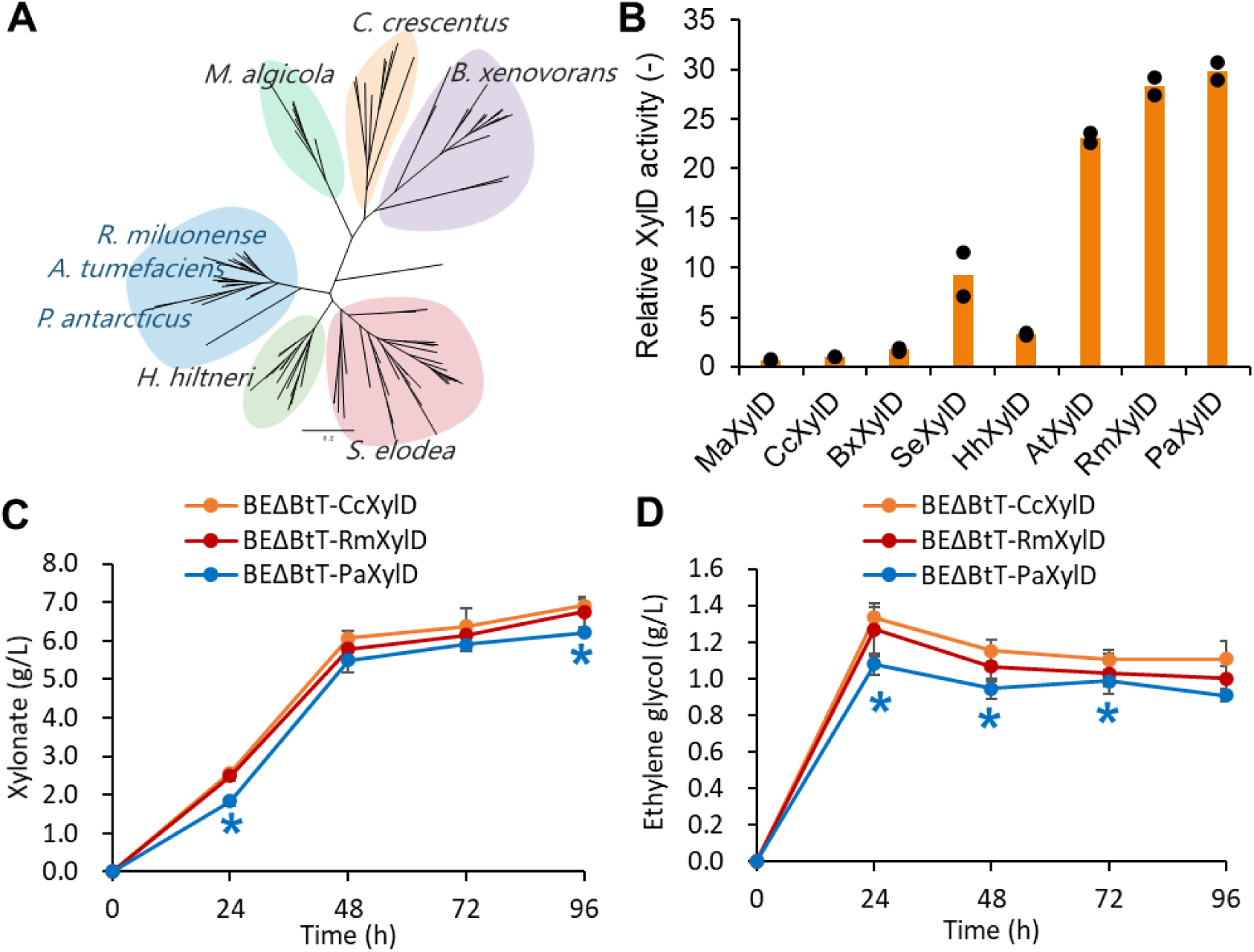
Effect of highly active XylD on xylonate accumulation and ethylene glycol production. (A) Phylogenetic tree analysis of XylD; (B) XylD activity of enzymes derived from species selected from the phylogenetic analysis. Orange bars show means of individual results (dots) obtained from two independent experiments; (C,D) Accumulation of xylonate and production of ethylene glycol during mixed sugar fermentation. Orange, red, and blue lines indicate the metabolite concentrations of BEΔBtT-CcXylD, BEΔBtT-RmXylD and BEΔBtT-PaXylD, respectively. Values are the mean ± SD of three biological replicates. Significant differences against BEΔBtT-CcXylD were evaluated by a two-tailed non-paired Student’s *t* test (* P < 0.05).

### XylX selection is crucial for xylose-dependent growth through the Weimberg pathway

Another promising xylose oxidative pathway is the Weimberg pathway. We explored the ability of engineered yeast strains harboring the Weimberg pathway to utilize xylose as their sole carbon source. To this end, we developed the BDΔBtT-ksaD-CcxylX strain by introducing XylX from *C. crescentus* and KsaD from *C. glutamicum* into the BDΔBtT strain, which is engineered for enhanced iron uptake. As shown in Figure 5A, neither the wild type YPH499 strain nor the BDΔBtT-ksaD-CcxylX strain showed any detectable growth on xylose.

**Figure 5.**
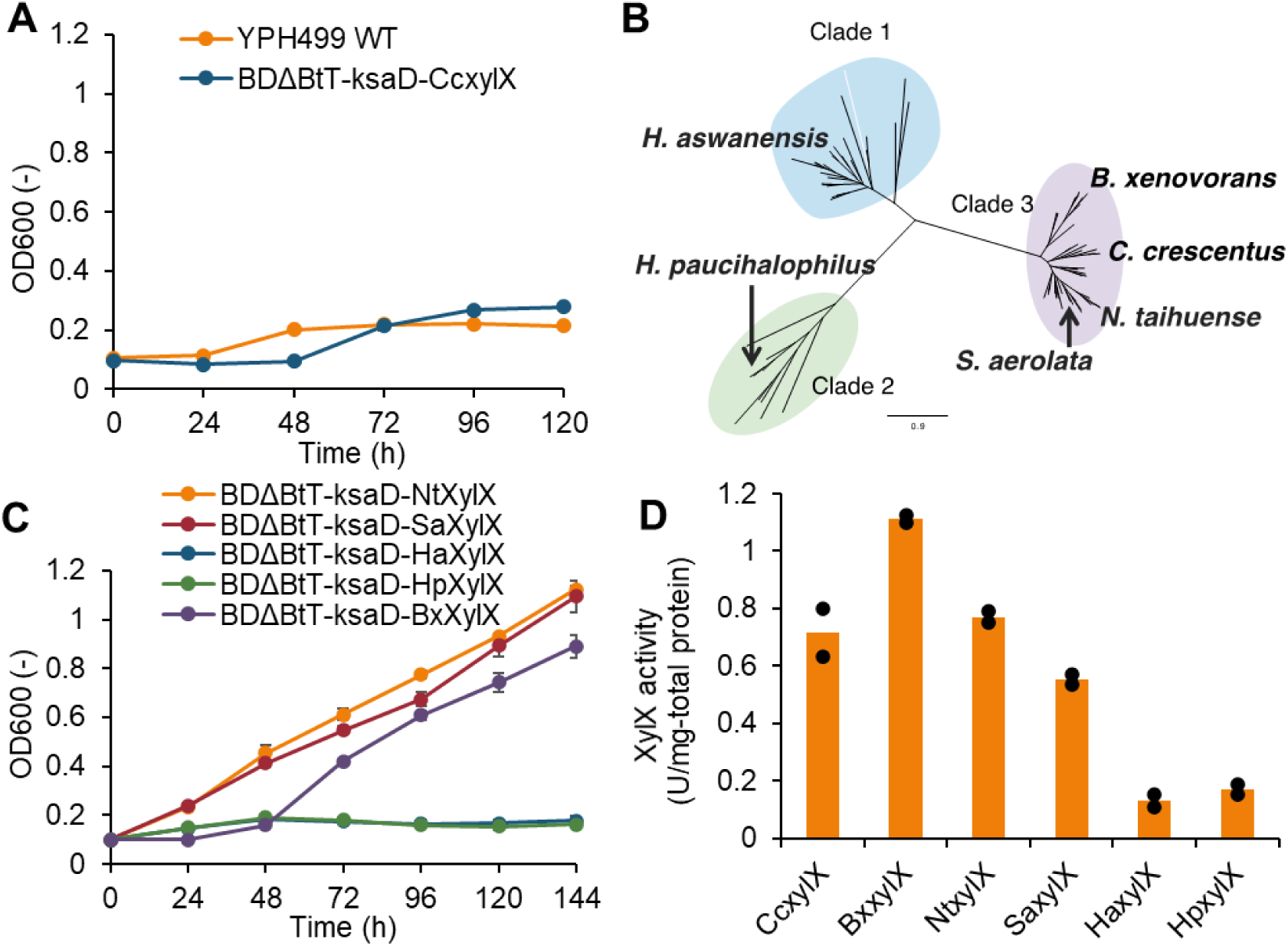
XylX screening to show xylose-dependent growth via Weimberg pathway. (A) Results of cell growth test on xylose via the Weimberg pathway (Orange: YPH499 WT, Red: BDΔBtT-ksaD-CcxylX, Blue: BDΔBtT-CcxylA-CcxylX). Growth tests were performed on medium containing 20 g/L of xylose as a sole carbon source. Values are the mean ± SD of three biological replicates; (B) Phylogenetic analysis of KDX dehydratase (XylX); (C) Introduction of *xylX* homologs increased cell growth on xylose. Values are the mean ± SD of three biological replicates; (D) KDX dehydratase activity of XylX homologs. Orange bars show the means of individual results (dots) obtained from two independent experiments.

Among the Weimberg pathway enzymes, efforts have been made to optimize XylB, XylD, XylA. As for xylose dehydrogenase, expression of *xylB* from *C. crescentus* has been demonstrated to be better suited for xylonate production than the NADP^+^-dependent dehydrogenase-encoding genes *xyd1* and *SUS2DD* (Toivari et al. 2010; Toivari et al. 2012). Wasserstrom et al. (2018) evaluated a total of four xylonate dehydratases isolated from *C. crescentus*, *Burkholderia cenocepacia*, *E. coli*, and *Ellin329*, even though yeast strains harboring these enzymes still accumulate xylonate. Likewise, xylonate dehydratases encoding *yjhG* from *E. coli* have been shown to be inefficient for the production of 1,2,4-butanetriol (Bamba et al. 2019). As an αKG semialdehyde dehydrogenase, expression of KsaD from *Corynebacterium glutamicum* had higher enzyme activity than XylA from *C. crescentus* (Borgström et al. 2019). Although the enzyme XylX from *C. crescentus* has a low reaction rate (kcat) and substrate specificity compared to other enzymes in the Weimberg pathway (Tai et al. 2016; Sutiono et al. 2020), optimization trials of XylX in yeast have been insufficient compared to the enzymes mentioned above.

Encouraged by the results obtained for XylD selection via phylogenetic tree analysis, we engineered yeast strains with alternative XylX enzymes identified using the same analytical method. The enzyme XylX was divided into three clades based on phylogenetic tree analysis (Figure 5B). Several representative XylX enzymes selected from each clade (*Halopiger aswanensis*, *Halalkalicoccus paucihalophilus*, *Burkholderia xenovorans*, *Sphingomonas aerolata*, and *Novosphingobium taihuense*) and introduced them into the BDΔBtT-ksaD strain harboring XylB and XylD from *C. crescentus* and KsaD from *C. glutamicum*. Subsequent growth assays on xylose, shown in Figure 5C, revealed that while the two XylXs from *H. aswanensis* and *H. paucihalophilus* did not support growth, the three XylXs from *B. xenovorans*, *S. aerolata*, and *N. taihuense* in Clade 3 enabled the growth on xylose upon replacement of the XylX from *C. crescentus*.

To investigate the relationship between xylose-dependent growth and XylX enzyme activity, we measured the KDX dehydratase activity of each enzyme (Figure 5D). As expected, XylXs from *H. aswanensis* and *H. paucihalophilus* showed significantly lower KDX dehydratase activities compared to those from Clade 3, suggesting that XylX from Clades 1 and 2 may not be suitable for constructing the Weimberg pathway in yeast. Conversely, among the enzymes belonging to Clade 3, XylX from *C. crescentus* was found to be inefficient for supporting xylose-dependent growth, despite its high KDX dehydratase activity (Figure 5A).

Next, our objective was to optimize XylA through phylogenetic tree analysis of all amino acid sequences registered as XylA in the NCBI database (Supplemental Fig. S1A). To assess the impact of XylA variants on xylose-dependent growth by engineered yeast, we introduced a representative XylA from each clade into the BDΔBtT-BxXylX strain, which harbors XylB and XylD from *C. crescentus*, and XylX from *B. xenovorans*. The XylA homologs tested were derived from *Asticcacaulis endophyticus*, *Streptomyces hygroscopicus*, *Pseudomonas inefficax*, *Bacteroidetes bacterium*, *Rhodopirellula sp. TMED11*, *Pseudomonas fluorescens*, *Sphingomonas aurantiaca*, *Rhodobacteraceae bacterium*, *Burkhorderia xenoborans*, *Nitrincola lacisaponensis* and *Burkholderia lata (Burkholderia aenigmatica)*. We observed differences among the XylA homologs in terms of supporting growth on xylose, and identified KsaD, previously used as a XylA homolog (Borgström et al. 2019), as one of the most suitable enzymes (Supplemental Fig. S1B).

### Utilization of xylose via the xylose oxidative pathway promotes growth in a mixed sugar medium

We investigated the effects of XylX homologs on the growth of engineered yeast with the Weimberg pathway when a mixture of glucose and xylose is used. The engineered yeast strain incapable of growing on xylose only, which harbored XylX from *H. aswanensis* and *H. paucihalophilus*, exhibited lower cell densities in the mixed medium compared to a control strain (YPH499ΔGRE3). In contrast, the engineered yeast containing XylX from *B. xenovorans*, *S. aerolata*, and *N. taihuense* exhibited higher cell densities than a control strain expressing XylX from *C. crescentus* (Figure 6A). When the xylose oxidative pathway is introduced in *S. cerevisiae*, the conversion of xylonate to KDX becomes a rate-limiting step, leading to the accumulation of a large amount of xylonate in the medium. Strains that demonstrated growth enhancement due to the replacement of selected enzymes via phylogenetic tree analysis accumulated less xylonate than a parental strain expressing enzymes from *C. crescentus* (Figure 6B). Therefore, our findings suggest that the increase in growth capability in the mixed sugar conditions is due to enhanced enzyme activities in the Weimberg pathway. Nevertheless, even in the xylose-utilizing strains, between 48.0% to 48.9% of the supplemented xylose remained unconverted and accumulated in the medium as xylonate.

**Figure 6.**
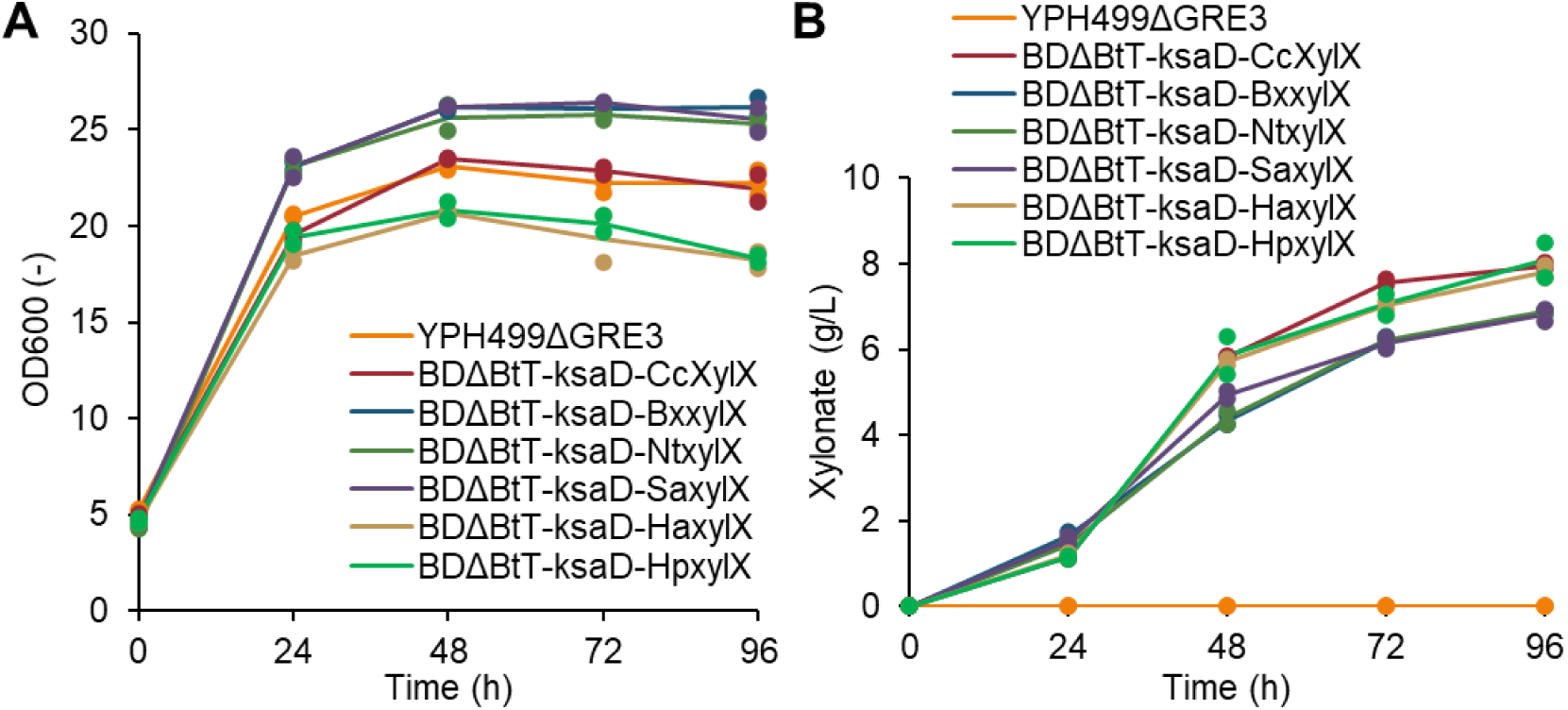
Cell growth and xylonate accumulation in mixed sugar culture by Weimberg pathway strains. Yeast culture tests were performed on a medium containing 10 g/L glucose and 10 g/L xylose. Line graphs show the means of individual results (dots) obtained from two independent experiments.

## Discussion

In *S. cerevisiae*, the oxidative xylose metabolic pathway has been used to produce 1,2,4-butanetriol, 3,4-dihydroxybutyric acid, ethylene glycol, and glycolic acid (Salusjärvi et al. 2017; Bamba et al. 2019; Yukawa et al. 2023). The enhancement of Fe uptake through *BOL2* disruption and t*TYW1* overexpression in this study may further improve the efficiency of these production processes. Although a higher titer of ethylene glycol has been achieved via the L-xylulose-1-phosphate pathway in *S. cerevisiae* (Uranukul et al. 2019), considerable room remains for improving titers via the Dahms pathway, as 80% of xylose was accumulated as xylonate (Figure 3). Despite the application of the phylogenetic enzyme search in this study, further enhancement of XylD activity is required. Given the resistance to fermentation inhibitors of lignocellulose degradation products and low pH, improving the xylose oxidative pathway in yeast could potentially be used to produce a variety of useful substances on an industrial scale.

Several studies have investigated the utility of enzymes from sources other than *C. crescentus* for various applications. These include using xylose dehydrogenase xyd1 from *Trichoderma reesei* for xylonate production, KDX aldolase *yagE* from *E. coli* for glycolate production via the Dahms pathway (Salusjärvi et al. 2017), and *ksaD* from *C. glutamicum* for constructing the Weimberg pathway (Borgström et al. 2019). XylD, in particular, exhibits low activity and becomes a bottleneck in the oxidative metabolism of xylose in yeast. For example, the xylonate dehydratase from *E. coli*, YjhG, was found to be inactive in *S. cerevisiae* (Wasserstrom et al. 2018), indicating a need to explore a wider range of potential organisms as potential enzyme sources for improving xylose oxidative metabolism. In this study, phylogenetic tree analysis was used to identify enzymes that are highly active in xylose oxidative metabolism. By introducing these enzymes, we have, to the best of our knowledge, enabled *S. cerevisiae* to grow on xylose as its sole carbon source via both the Dahms and Weimberg pathways for the first time (Figures 2 and 5). Thus, the search for, and identification of, highly active enzymes represent a powerful approach for efficiently constructing heterologous pathways in budding yeast.

XylXs from *B. xenovorans*, *S. aerolata*, and *N. taihuense*, classified within Clade 3, facilitate cell growth on xylose as the sole carbon source via the Weimberg pathway. Conversely, despite the similarity in activity of XylX from *C. crescentus* and that from *N. taihuense*, engineered strains expressing XylX from *C. crescentus* did not grow on xylose (Figure 5). This finding is consistent with earlier reports (Borgström et al. 2019; Wasserstrom et al. 2018). Previous studies have suggested that the activity of KDX dehydratase may be inhibited by xylonate, an intermediate metabolite of the Weimberg pathway. When constructing oxidative xylose metabolism pathways using *S. cerevisiae* as a host, the low activities of XylD result in substantial accumulation of xylonate both inside and outside the cell (Salusjärvi et al. 2017; Shen et al. 2020). Consequently, it is suggested that the inhibition of CcXylX by high intracellular concentrations of xylonate in *S. cerevisiae* could explain why xylose cannot be utilized for growth.

## Author contributions

TY, TB and TH conceived and designed the study. TY, MW and RK conducted experiments. TK, TY, TB and MW analyzed the data. All authors discussed the results. TK and TY wrote the manuscript. All the authors have read and approved the final manuscript. TH supervised all aspects of this study.

## Data availability

The data obtained and/or analyzed in this study are available from the corresponding author upon reasonable request.

## Declarations

### Ethical approval

This article does not contain any studies involving human participants or animals.

