## Supplemental information for "Engineering *Saccharomyces cerevisiae* for growth on xylose using an oxidative pathway"

**Supplemental Figure**

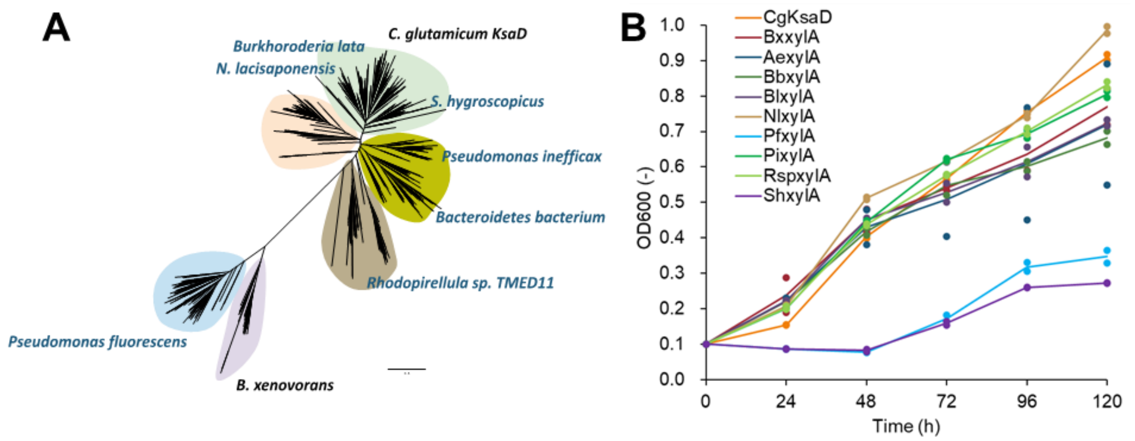


**Supplemental Fig. 1 XylA screening based on phylogenetic tree analysis.** (A) Phylogenetic tree analysis of XylA; (B) Yeast growth tests were performed on medium containing 20 g/L of xylose as the sole carbon source. The line graph shows the means of individual results (dots) obtained from two independent experiments.

**Supplemental Table 1. Plasmids used in this study**

| Plasmid name | Description |
|---|---|
| pTS-A-xylBD | pTDH3-xylB, pSED1-xylD, ADE2 marker |
| pIAur-tTYW1 | PGK1p-tTYW1-PGK1t, AUR1-C marker |
| pIL-pTDH3 tADH1 | pTDH3-tADH1, LEU2 marker |
| pIL-pTDH3-yagE | pTDH3-yagE-tADH1, LEU2 marker |
| pIU-pTDH3-CgksaD | pTDH3-CgksaD-tADH1, URA3 marker |
| pIL-pSED1-tSAG1 | pSED1p-tSAG1, LEU2 marker |
| pIL-pSED1-HaxylX-tSAG1 | pSED1p-HaxylX-tSAG1, LEU2 marker |
| pIL-pSED1-HpxylX-tSAG1 | pSED1p-HpxylX-tSAG1, LEU2 marker |
| pIL-pSED1-CcxylX-tSAG1 | pSED1p-CcxylX-tSAG1, LEU2 marker |
| pIL-pSED1-BxxylX-tSAG1 | pSED1p-BxxylX-tSAG1, LEU2 marker |
| pIL-pSED1-NtxylX-tSAG1 | pSED1p-NtxylX-tSAG1, LEU2 marker |
| pIL-pSED1-SaxylX-tSAG1 | pSED1p-SaxylX-tSAG1, LEU2 marker |
| pIA-pTDH3-tADH1 | pTDH3-tADH1, ADE2 marker |
| pIA-pTDH3-xylB | pTDH3-xylB-tADH1, ADE2 marker |
| pGK406 | Empty vector, URA3 marker |
| pIU-pTDH3-tADH1 | pTDH3-tADH1, URA3 marker |
| pIU-pTDH3-BxxylA | pTDH3-BxxylA-tADH1, URA3 marker |
| pIU-pTDH3-AexylA | pTDH3-AexylA-tADH1, URA3 marker |
| pIU-pTDH3-ShxylA | pTDH3-ShxylA-tADH1, URA3 marker |
| pIU-pTDH3-PixylA | pTDH3-PixylA-tADH1, URA3 marker |
| pIU-pTDH3-BbxylA | pTDH3-BbxylA-tADH1, URA3 marker |
| pIU-pTDH3-RspxylA | pTDH3-RspxylA-tADH1, URA3 marker |
| pIU-pTDH3-PfxylA | pTDH3-PfxylA-tADH1, URA3 marker |
| pIU-pTDH3-SaxylA | pTDH3-SaxylA-tADH1, URA3 marker |
| pIU-pTDH3-RbxylA | pTDH3-RbxylA-tADH1, URA3 marker |
| pIU-pTDH3-BlxylA | pTDH3-BlxylA-tADH1, URA3 marker |
| pIU-pSED1-tSAG1 | pSED1p-tSAG1, URA3 marker |
| pIU-pSED1-MaxylD | pSED1p-MaxylD-tSAG1, URA3 marker |
| pIU-pSED1-CcxylD | pSED1p-CcxylD-tSAG1, URA3 marker |
| pIU-pSED1-BxxylD | pSED1p-BxxylD-tSAG1, URA3 marker |
| pIU-pSED1-HexylD | pSED1p-HexylD-tSAG1, URA3 marker |
| pIU-pSED1-SexylD | pSED1p-SexylD-tSAG1, URA3 marker |

| | |
|---|---|
| pIU-pSED1-AtxylD | pSED1p-AtxylD-tSAG1, URA3 marker |
| pIU-pSED1-RmxylD | pSED1p-RmxylD-tSAG1, URA3 marker |
| pIU-pSED1-PaxylD | pSED1p-PaxylD-tSAG1, URA3 marker |

**Supplemental Table 2. Primers used in this study**

| Primer | Sequence |
|---|---|
| xhoI-SED1p F | cgggccccccctcgagattggatatagaaaattaacgtaagg |
| xhoI-SED1p R | aactgtacacccgggcttaatagagcgaacgtattttattttg |
| SAG1t F | cccgggtgtacagttagtacattgagtc |
| SAG1t R | accgcggtggcggccgcatccagtgagcgcgcgtaatacgac |
| dBOL2_ ADE2 F | ttcttgaataatacataacttttc |
| dBOL2_ ADE2 R | ataagtgatcttatgtatgaaattc |
| BOL2 up F | acgttctctccgttgttcaaacccc |
| BOL2 up R | agttatgtattattcaagaaatatatgtatatatataacaccg |
| BOL2 down F | tcatacataagatcacttataaaggatgatattgttctattattaag |
| BOL2 down R | acagcaacgacgacaatgccaaacc |

**Supplemental Table 4. Yeast strains constructed in this study**

| Strain name | Description |
|---|---|
| YPH499 | MATa ura3-52 lys2-801_amber ade2-101_ochre trp1-Δ63 his3-Δ200 leu2-Δ1 |
| YPH499ΔGRE3 | YPH499, gre3Δ::kanMX4 |
| BD | YPH499ΔGRE3, pTS-A-xylBD |
| BDE | BD, pIL-TDH3p-yagE |
| BDΔB-E | BD, bol2Δ::HIS3, pIL-TDH3p-yagE |
| BDΔBtT | BD, bol2Δ::HIS3, pIAur-tTYW1 |
| BDΔBtT-E | BDΔBtT, pIL-TDH3p-yagE |
| BDΔBtT-H | BDΔBtT, pIL-TDH3p-yagE |
| BE | YPH499ΔGRE3, pIA-pTDH3-xylB, pIL-TDH3p-yagE |
| BEΔBtT | BE, bol2Δ::HIS3, pIAur-tTYW1 |
| BEΔBtT-MaxylD | BEΔBtT, pIU-pSED1-MaxylD |
| BEΔBtT-CcxylD | BEΔBtT, pIU-pSED1-CcxylD |
| BEΔBtT-BxxylD | BEΔBtT, pIU-pSED1-BxxylD |
| BEΔBtT-HexylD | BEΔBtT, pIU-pSED1-HexylD |
| BEΔBtT-SexylD | BEΔBtT, pIU-pSED1-SexylD |
| BEΔBtT-AtxylD | BEΔBtT, pIU-pSED1-AtxylD |
| BEΔBtT-RmxylD | BEΔBtT, pIU-pSED1-RmxylD |
| BEΔBtT-PaxylD | BEΔBtT, pIU-pSED1-PaxylD |
| BDΔBtT-ksaD | BDΔBtT, pIU-pTDH3-CgksaD |
| BDΔBtT-ksaD-CcxylX | BDΔBtT-ksaD, pIL-pSED1-CcxylX-tSAG1 |
| BDΔBtT-ksaD-HaxylX | BDΔBtT-ksaD, pIL-pSED1-HaxylX-tSAG1 |
| BDΔBtT-ksaD-HpxylX | BDΔBtT-ksaD, pIL-pSED1-HpxylX-tSAG1 |
| BDΔBtT-ksaD-BxxylX | BDΔBtT-ksaD, pIL-pSED1-BxxylX-tSAG1 |
| BDΔBtT-ksaD-SaxylX | BDΔBtT-ksaD, pIL-pSED1-SaxylX-tSAG1 |
| BDΔBtT-ksaD-NtxylX | BDΔBtT-ksaD, pIL-pSED1-NtxylX-tSAG1 |
| BDΔBtT-BxxylX | BDΔBtT, pIL-pSED1-BxxylX-tSAG1 |
| BDΔBtT-BxxylX-BxxylA | BDΔBtT-BxxylX, pIU-pTDH3-BxxylA |
| BDΔBtT-BxxylX-AexylA | BDΔBtT-BxxylX, pIU-pTDH3-AexylA |
| BDΔBtT-BxxylX-ShxylA | BDΔBtT-BxxylX, pIU-pTDH3-ShxylA |
| BDΔBtT-BxxylX-PixylA | BDΔBtT-BxxylX, pIU-pTDH3-PixylA |
| BDΔBtT-BxxylX-BbxylA | BDΔBtT-BxxylX, pIU-pTDH3-BbxylA |

| | |
|---|---|
| BDΔBtT-BxxylX-RspxylA | BDΔBtT-BxxylX, pIU-pTDH3-RspxylA |
| BDΔBtT-BxxylX-PfxylA | BDΔBtT-BxxylX, pIU-pTDH3-PfxylA |
| BDΔBtT-BxxylX-SaxylA | BDΔBtT-BxxylX, pIU-pTDH3-SaxylA |
| BDΔBtT-BxxylX-RbxylA | BDΔBtT-BxxylX, pIU-pTDH3-RbxylA |
| BDΔBtT-BxxylX-BlxylA | BDΔBtT-BxxylX, pIU-pTDH3-BlxylA |
